# Investigating polarity effects in DNA base stacking

**DOI:** 10.1101/2025.09.25.678614

**Authors:** Jibin Abraham Punnoose, Chai S. Kam, Tristan Melfi, Sweta Vangaveti, Alan A. Chen, Ken Halvorsen

## Abstract

Nucleic acid structures are stabilized by both base pairing and base stacking. While energetics of base pairing interactions are relatively well established, our understanding of the energetic contributions of base stacking remain incomplete. Here, we use a combination of single-molecule and computational biophysics approaches to investigate the effect of strand polarity on base-stacking energetics. We designed pairs of DNA constructs with reversed stacking polarities at nick sites, along with corresponding no-stack controls to isolate stacking contributions. Performing single-molecule force-clamp assays with a Centrifuge Force Microscope (CFM), we observed polarity-dependent differences in stacking energetics. These differences were most pronounced in purine–purine and certain purine–pyrimidine interactions. Notably, a 5′ purine stacked on a 3′ pyrimidine was generally more stable than the reverse polarity. We employed molecular dynamics (MD) simulations to observe stacking interfaces in the DNA constructs. The simulations were qualitatively consistent with our experiments, and showed positional differences between opposite polarity stacking pairs, giving some insight into the origin of these polarity differences. Overall, these results demonstrate that base polarity can modulate stacking stability and should be considered when designing short duplex regions such as overhangs in molecular biology and biotechnology applications.

## Introduction

Base stacking is a critical non-covalent interaction between aromatic ring structures in DNA and RNA bases that plays an important role in stabilizing simple and complex nucleic acid structures. Stacking between adjacent nucleobases provides van der Waals interactions and hydrophobic effects that reinforce duplex formation and maintain the integrity of secondary and tertiary structures such as hairpins, pseudoknots, and kissing loops [1-3]. Base stacking also plays key roles in processes such as strand invasion and displacement [4,5], the assembly and resilience of nucleic acid nanomaterials [6-10], and the efficiency of biochemical reactions including DNA ligation [11-13]. Various experimental methods have been used to quantify stacking energetics, including electrophoretic mobility analysis of nicked duplexes [14,15], optical tweezers pulling stacked DNA nanostructures [16], single-molecule force clamps applied to nicked duplexes [12,17], kinetic analysis of single-molecule fluorescence [18], and thermal melting analysis [19]. The most recent of these have enabled quantification of stacking interactions between two individual bases. Across these approaches, base-stacking energies in DNA have been found to range from ∼-0.5 to ∼-3 kcal/mol, with purine–purine stacks typically being the most stable and pyrimidine–pyrimidine the least.

Despite substantial recent progress in quantifying stacking interactions, the role of base polarity (i.e. the 5′–3′ directionality of stacked nucleotides) has not been systematically investigated (**Figure 1**). In our own lab’s recent work, we began with the assumption that polarity would be unlikely to have a major effect on stacking energetics, and focused on the 10 unique dinucleotide combinations [12]. More recent studies suggested polarity may affect base stacking energies [18], including our own follow up work on base stacking in DNA nanostructures [10]. In that work, we found that in a DNA tetrahedron, polarity of the terminal stacking interactions on the joining sticky ends can influence nanostructure stability. This was particularly true in purine–pyrimidine pairs, where a pyrimidine such as cytosine in the 5′ position relative to a purine resulted in decreased stability across nicked interfaces in DNA tetrahedra [10].

**Figure 1.**
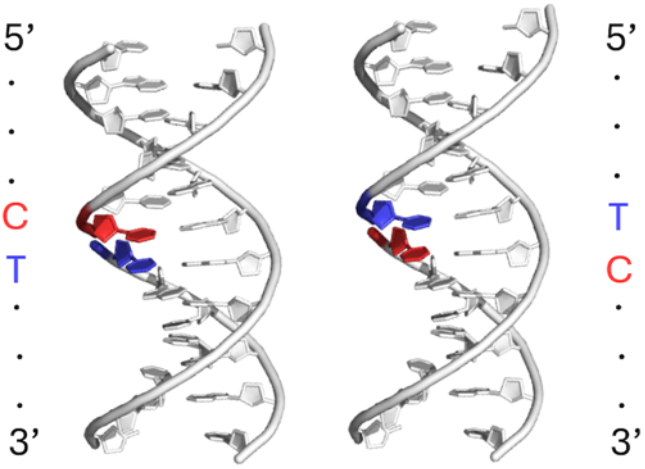
Illustration of a duplex with a cytosine and thymine at a nicked interface. The nicked strand is 5’ at the top and 3’ at the bottom. The base stacking interaction across the nick would be denoted as C|T (left) and T|C (right).

To investigate this effect in a controlled and quantitative manner, we employed a single-molecule force-clamp assay using the centrifuge force microscope (CFM) [20]. The CFM enables high-throughput application of centrifugal force to DNA tethers while monitoring microsphere displacement in real time [21-23]. For this study, we designed DNA constructs that differ only in the polarity of the stacked bases across a nick (either 5′ purine stacked on 3′ pyrimidine or vice versa) compared with analagous stacking experiments we previously completed for a subset of pairs [12]. This design isolates the base-stacking contribution and enables precise measurement of polarity-dependent energetic differences. To complement the single-molecule measurements, we performed Molecular Dynamic (MD) simulations of representative models of the same constructs at different temperatures. Analysis of the three-dimensional positions and movements of the interfacial stacking bases allowed further exploration of the reasons behind the difference in the polarity of base stacks.

Our results demonstrate and quantify measurable differences in stacking energetics between depending on the polarity of stacked bases. We observed the largest polarity effects in purine–purine and some purine–pyrimidine combinations, up to -1.2 kcal/mol between G|C stack and the weaker C|G stack. Our MD results qualitatively reproduce the stronger stacking polarity, and the detailed positions and motions of the stacking bases provide insights into the molecular mechanisms underlying these polarity differences. Overall, our findings highlight stacking polarity as an important and previously underappreciated factor influencing nucleic acid thermodynamics and underscore the need to consider base orientation when designing short duplex features—such as overhangs or nicked regions—in molecular biology and nanotechnology applications.

## Materials and Methods

### DNA and oligonucleotides

The sequences of all DNA oligonucleotides used in this study are listed in **Table S1**. Oligonucleotides were obtained from Integrated DNA Technologies and were PAGE-purified (www.idtdna.com). M13mp18 circular single-stranded DNA (New England Biolabs, catalog # N4040S) served as the scaffold for construct synthesis.

### Construct Preparation

The circular single-stranded M13mp18 plasmid was linearized based on previously established protocol [12]. Briefly, 5 μL of M13mp18 (∼100 nM) was mixed with 2.5 μL of 10x CutSmart buffer (NEB, catalog # B6004S), 1 μL of a 100 μM oligonucleotide containing the BtsCI recognition site, and 16.5 μL of nuclease-free water in a PCR tube. The mixture was heated to 95 °C for 30 seconds to denature secondary structures and then cooled to 50 °C. At this temperature, the BtsCI enzyme (New England Biolabs, catalog # R0647S) was added and the reaction was incubated for 15 minutes to allow site-specific nicking. The enzyme was subsequently inactivated by heating to 95 °C for 1 minute, followed by cooling to 4 °C.

DNA constructs were then assembled by hybridizing 124 staple oligos to the linearized M13 scaffold, as described previously [12]. The staple strand annealing to the 3′ end of the scaffold had a 5′ dual biotin for immobilization to streptavidin-coated surfaces or beads. The staple at the 5′ end extended beyond the scaffold sequence, forming a 30-nucleotide 3′ single-stranded overhang. A complementary oligo, either 38 or 41 nucleotides in length, with 30 nucleotides matching the 5′ overhang and an additional 8–11 base extension, was used to form the duplex tether between the glass surface and microsphere (**Figure 2a, Table S2**). Construct variants were generated by altering only the oligo hybridizing to the 5′ end of the scaffold (**Table S3**).

**Figure 2:**
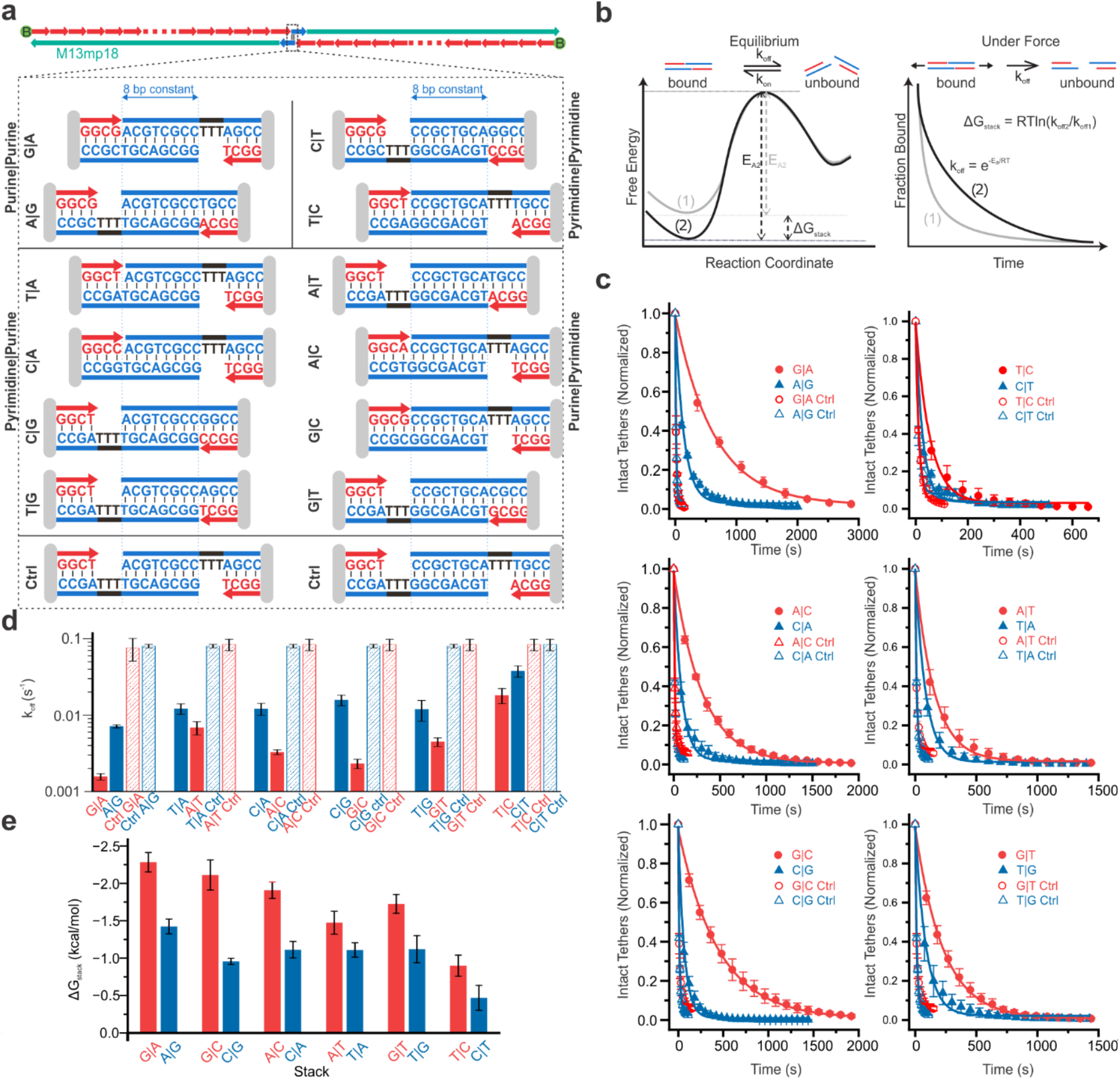
Polarity dependence of base stacking quantified by single-molecule force-clamp assay. (a) DNA constructs were assembled on linearized M13mp18 ssDNA (green strand) using 123 oligos (red strands). One end oligo contains a double biotin (green spheres) for surface immobilization. The other end oligo extends beyond the M13 ssDNA to provide a platform for a “programmable” oligo (blue strand) bearing 8 or 11 nucleotide overhangs that define the terminal base pair and base stack. Control constructs retain the same base pairs but lack terminal stacking. (b) Free energy diagram under equilibrium showing an increase in activation energy (ΔG_stack_) due to base stacking. Under force, the activation barrier is lowered and rebinding is suppressed, allowing exponential dissociation suitable for kinetic analysis. (c) Dissociation curves and single-exponential fits measured at 15 pN for stacked constructs (closed symbols) and their controls (open symbols), based on experimental replicates pooled randomly into three groups. (d) Measured off-rates (k_off_) for various base stacks and controls obtained from single-exponential fitting of decay curves. Error bar represents standard deviation of the off-rates obtained from three experimental groups (e) Base stacking energies calculated from the off-rates. Error bar represents propagated error from the off-rates.

### DNA construct Immobilization on the magnetic beads

To attach DNA constructs to magnetic microspheres, 20 μL of streptavidin-coated Dynabeads M-270 (2.8 μm diameter, Thermo Fisher Scientific, catalog #65306) were washed three times with 100 μL of 0.1% Tween-20 in 1× phosphate-buffered saline (PBS). The final pellet was resuspended in 10 μL, to which 10 μL of ∼500 pM DNA construct was added. The mixture was vortexed at 1000 rpm for 20 minutes to promote binding. Unbound DNA was removed by three additional washes with 100 μL of PBST (PBS + 0.1% Tween-20), and the final bead suspension was adjusted to 40 μL.

### Chamber Preparation and construct immobilization on the surface

Microfluidic chambers for single-molecule force measurements were assembled as previously described [12,23]. Each chamber was formed by sandwiching two parallel strips of ∼2 mm wide Kapton tape (Bertech, cat. no. PPTDE-1/4) between an 18-mm and a 12-mm circular glass coverslip (Electron Microscopy Sciences, catalog #72230-01 and #72222-01). The assembly was mounted on an SM1A6-threaded adapter (ThorLabs, Newton, NJ) using the Kapton tape. The surface was functionalized by introducing 5 μL of 0.1 mg/mL streptavidin in 1× PBS and incubating for 1 minute, followed by three PBST washes to remove unbound protein. Next, 5 μL of ∼500 pM biotinylated DNA construct was introduced and incubated for 5 minutes to allow immobilization on the glass surface. Excess unbound constructs were removed by washing with PBST. DNA-coated microspheres were then flowed into the chamber, and the chamber was sealed using vacuum grease. The reaction chamber mounted on the SM1A6 adaptor was then screwed into an assembly consisting of SM1-05 inch tube. The chamber was incubated in an inverted orientation for 10 minutes to allow complementary strands to hybridize and tether between the glass surface and the microspheres. Immediately after incubation, the chamber assembly was mounted into the CFM for measurements.

### Force Clamp Assay

The sealed sample chamber was placed into the CFM module and loaded into a Sorvall Legend X1R benchtop centrifuge equipped with TX-400 rotor and 400 mL Swinging buckets. A counterbalance of equal mass and center of gravity was used to ensure rotational stability. Instrument control, including rotation speed and camera settings, was achieved via custom-written LabVIEW software. For these experiments, image acquisition occurred at 2 frames per second (fps), while data were saved at a reduced rate of 0.2 fps to manage file size during long-duration measurements (typically up to 1 hour).

The force exerted on the DNA tethers was calculated as the centrifugal force on the microspheres, using the expression F=mω^2^r, where m is the effective mass of the bead (accounting for buoyancy), ω is the angular velocity, and r is the radial distance from the rotor axis to the sample chamber (measured as 0.119 m). Based on prior calibration, the effective mass of Dynabeads M-270 was estimated to be 6.9 × 10^−12^ g [12]. For the experiments reported here, we applied 1291 RPM, corresponding to a force of ∼15 pN, and monitored the system until >95% of tethers had dissociated. The time point corresponding to the first frame after reaching the target RPM was defined as time zero for dissociation analysis.

### Off-rate measurement & free energy calculation

Image analysis for the CFM experiments was carried out using a custom MATLAB script, previously developed in our lab [12,23]. The script employs the “imfindcircles” function to automatically identify circular features corresponding to DNA-tethered microspheres. All automatically identified features were manually screened to confirm that they represented valid, individual tethers. Non-circular particles, aggregated beads, debris, and out-of-focus objects were excluded from further analysis. Beads that appeared irregular or ambiguous—such as those closely clustered or exhibiting non-spherical geometry—were also excluded to minimize the likelihood of analyzing multiple tethers. Following bead identification in the initial frame, the program tracked the variance in pixel intensity at each bead’s location across the full duration of the image sequence. Dissociation events were identified by a marked drop in intensity variance—corresponding to the abrupt transition from high contrast (bound state) to low contrast (unbound state). Beads that showed multiple-step dissociation profiles were rare due to rigorous pre-screening and were excluded from the analysis to avoid contributions from multiple tethering events.

Dissociation time data were compiled and plotted as histograms, using bin widths adjusted to ensure a consistent number of bins across datasets, even when the overall experiment duration varied by more than an order of magnitude. Decay curves were fit in OriginLab using a single-exponential model: y = y_0_ + A × e^−kt,^ where y is the fraction of tethers remaining at a given time t, y_0_ is the y-axis offset or the baseline, A is the fraction of tethers at the beginning of the experiment (typically 1) and k is the off-rate for that particular force.

Each experimental condition was replicated at least three times, and dissociation rate constants (k) were determined individually for each replicate. The reported off-rates represent the mean and standard deviation of the replicate values. Base-stacking free energies were extracted by comparing the dissociation rates of the stacking construct to those of a corresponding control lacking the base stack, assuming an Arrhenius-type dependence of the off-rate on activation energy:

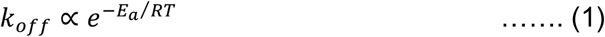

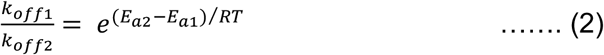

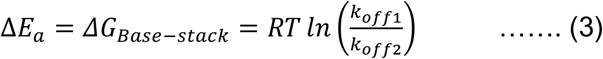

In these expressions, k_off1_ and k_off2_ represent the off-rates for the constructs with and without the base stack, respectively; E_a1_ and E_a2_ are the corresponding activation energies; and ΔG_base-stack_ is the effective free energy contribution from the base-stacking interaction.

### Molecular dynamics simulations

Models of the nicked duplexes were created using Molecular Operating Environment (MOE) (2022.2; Chemical Computing Group) and simulated using openmm [24] version 8.2 with the tumuc1 force field. To better emulate real-world conditions, we applied position restraints on the terminal nucleotides to prevent fraying as these duplexes would be a part of larger nanostructures. And additionally, we removed the bond between the bases of interest as the modified “nicked” topology better represents experimental melting. Each simulation was done using the tumuc1 force field [25] and solvated using an explicit TIP3P water model. An additional neutralizing concentration of K+ ions [27] was also added. To further explore the effect of base stacking on thermal stability each duplex was simulated in an increasing temperature series. With simulations done at 300k – 400k in increments of 20k. Altogether, each duplex was simulated 6 times with each simulation run for 250 nanoseconds. Positions were saved every 25 picoseconds, generating 10,000 data points per simulation.

Each simulation frame exists in a high dimensional space that requires projections into a lower dimension coordinate system for analysis. This new coordinate system must be robust enough to capture the behavior of the nicked interface while being simple enough interpretation. We described a 2-dimensional coordinate system (ρ, θ) to describe the relative conformations between the 2 nucleotides in the nicked interface. Coordinates are described about the geometric center of the 5’ nucleotide, 6 membered rings. The coordinate, ρ, is computed as the Euclidean distance between both centers. Describing how far apart each base is from one another. While θ, is computed from the angle formed between 2 vectors projected in the xy-plane. Specifically, the vectors that bisect the carbon atoms along the Watson-crick face on the range [0, 180] using the geometric definition of the dot product. When the nucleotides twist in the xy plane the angle between these 2 vectors track that twist.

Computations were performed using resources provided by the UAlbany AI+ initiative. Utilizing NVIDIA, A100 GPU’s providing quick turnaround in simulation time and analysis. Reaching simulation speeds of up to 1 microsecond per day.

## Results

Base-stacking interactions are relatively weak, and the hybridization energy of nucleic acids is highly sequence-dependent, making it difficult to measure stacking energies. Previously, we developed a framework for measuring these interactions that involves applying a single-molecule force clamp to DNA tethers held together with a short duplex. By altering the design to control for terminal base stacking, we can determine the energetic contribution of the individual stack from the unbinding kinetics of tethers with and without the terminal stack [12].

Here, the objective is to compare similar base stacks with different strand polarities. To overcome this challenge, we designed DNA tethers with short duplexes containing identical base pairs but differing in the presence or absence of a terminal base stack, enabling us to study all 12 base-stacking combinations (**Figure 2a**). A 3-nucleotide poly-T spacer was introduced in the control constructs to disrupt terminal stacking. This design isolates the contribution of a single base-stacking interaction between two bases, and allows direct comparison of stacking energies for opposite polarities. In our previous study [12], we quantified the stacking interactions of G|A, A|T, A|C, G|C, G|T, and C|T. In the present work, we extend this analysis to the corresponding opposite-polarity stacks A|G, T|A, C|A, C|G, T|G, and T|C.

We used a custom-built centrifuge force microscope (CFM) to perform high-throughput force-clamp assays [23]. In this setup, centrifugal force applies constant tension to microspheres tethered to a surface via DNA constructs, enabling the measurement of bond lifetimes and dissociation kinetics under controlled force (**Figure 2b**). Images of the microspheres under tension were captured using a machine vision camera coupled to a 40× objective, and transmitted wirelessly to an external computer. As the centrifuge spins, applied force leads to the dissociation of tethers, causing the corresponding microspheres to disappear from the field of view. Each microsphere is monitored individually to detect dissociation events, which are compiled into decay curves used to extract molecular off-rates (**Figure 2c & Figure S1-S3**).

We calculated stacking energies by comparing the off-rates of each base-stacked construct against its control (**Figure 2d-e**). The measured stacking energies (in kcal/mol) are presented in **Table 1** below, along with those previously measured to complete the table. These results clearly indicate that base-stacking interactions are sensitive to strand polarity, particularly in purine–purine and certain purine–pyrimidine pairs. In purine–pyrimidine stacks, stability tended to be higher when the purine occupied the 5′ position. The most pronounced polarity effect was observed in the G|C vs C|G pair, while the smallest differences were seen for A|T vs T|A and C|T vs T|C.

**Table 1:**
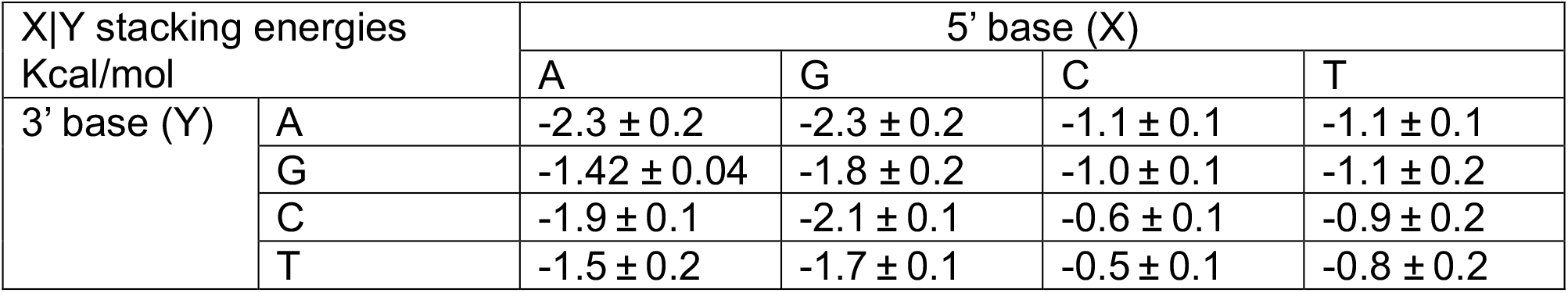
base-stacking energies in kcal/mol.

To further investigate the difference in base stacking polarities and ideally identify molecular causes of these differences, we turned to MD simulations. We generated three-dimensional atomistic models of nicked duplexes, comparable to those that we experimentally measured with single molecule experiments. Each system was solvated with water and a neutralizing amount of K+ ions. For each duplex, we ran six simulations of 250 nanoseconds each at increasing temperatures (from 300 K to 400 K) to observe how each nicked interface handles increasing temperatures (**Figure 3a**). We restrained the motion of the terminal ends of the duplex to prevent fraying to solely focus on the motions of the nicked interface. Each 250 nanoseconds simulation saved coordinates every 25 picoseconds resulting in 10,000 frames per simulation.

**Figure 3.**
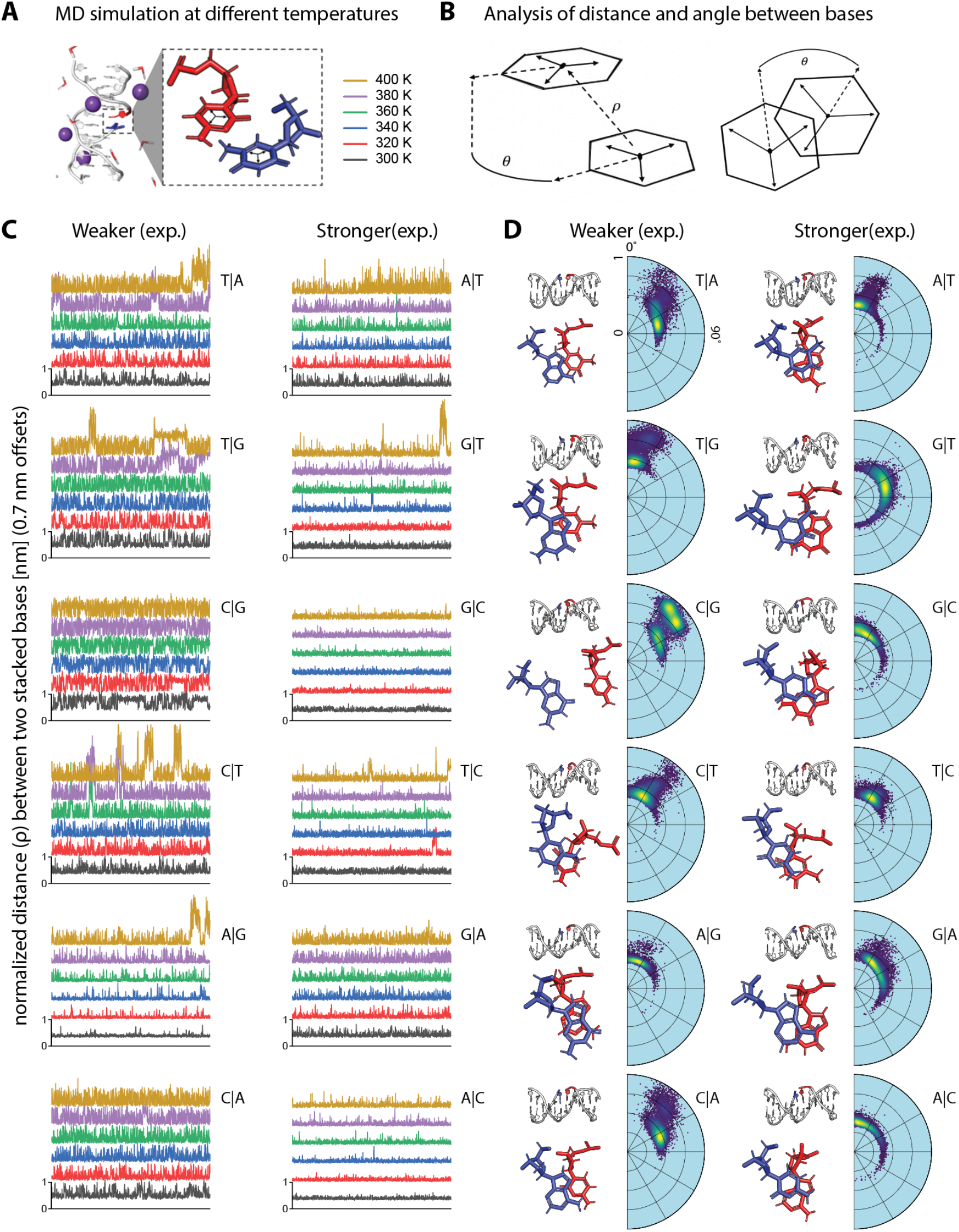
Molecular dynamics simulations of duplexes. a) Nicked duplexes were constructed and placed into an all-atom simulation at various temperatures. b) The nucleotides of interest are projected into a two-component coordinate system to bring the high dimensional motions of nucleic acids into a simplified representation of their dynamics. c) The distance (rho) between the two stacked bases across the nick is plotted for all conditions, with an offset between temperatures for visualization. d) The results of the 300 K simulations projected onto the polar plots containing the distance (rho) and the angle (theta), giving an overview of dynamics between the interfaces.

Collectively, these simulations result in an enormous amount of data that must be distilled to some useful parameters to enable comparison with experiments. We focused on the motion of the two nucleotides within the nicked interface, and defined a two component coordinate space. One component (rho) was defined by the Euclidean distance between the centers of the 6 membered rings, and the other (theta) was defined by an angular shift in direction of the Watson-crick edge of the bases (**Figure 3b**). We found that these two components are descriptive enough to capture the relative dynamics between both bases to then allow us to compare relative thermal stabilities.

First we examined the variation in the inter-base distance (rho), as functions of time and temperature (**Figure 3c**). For each polarity pair, we plotted the stronger ones from our experimental results on the right-hand side. It is obvious to see from this data that the stronger stacking results in a tighter distribution of rho values around a stable stacking distance of ∼ 0.5 nm at a given temperature. It can also be observed that experimentally weaker stacking pairs suffer more frequent large excursions from the stable stacking position at lower temperatures than their stronger counterparts. It can also be noted that the most striking visual differences between polarity pairs occur for those stacks that were experimentally measured to have the largest energy differences, notably A|C vs C|A and G|C vs C|G.

To further explore these interactions, we developed polar density plots for all conditions (**Figure S4**). For the temperature of 300 K where all stacks are expected to be relatively stable, the polar plots are paired with MD snapshots to illustrate the likely positions of the stacked pairs (**Figure 3d**). In these polar plots, the more favorable base stacks appear to dissipate thermal energy by rotating in the theta plane while maintaining that ∼0.5 nm distance between the pairs. Less favorable stacking interactions look more chaotic on the polar plots, with more frequent excursions outside of that ∼0.5 nm range as the alignment of partial charges is not as favorable. This information compliments the experimental results by independently supporting with observations on base stacking stability.

## Discussion

With this work, we provide direct experimental evidence that base stacking energy is affected by the strand polarity between the adjacent bases. In some cases such as G|C and C|G there is a ∼2 fold energy difference, while other cases like A|T and T|A are more subtle with differences on the order of 30%. With this data, we complete the quantitation of the single base stacking energetics of all 16 nucleotide combinations (**Table 1**). The polarity differences are largely supported by data from MD simulations, which shows more consistent localization between adjacent bases that were experimentally more energetically favorable. These simulations also show decreasing localization at increasing temperatures as the stacks become unstable.

For our experimental data, we can make some comparisons with other published studies. Our polarity differences qualitatively agree with another recent study performed using single-molecule fluorescence, but not quantitatively [18]. All of our polarity pairs agree between stronger and weaker, but there is some deviation in the exact values. Another single-molecule study quantified stacked pairs in blunt end duplexes [16], and similarly found G|C stronger than C|G and A|T stronger than T|A, but again the quantification had some deviation. As we previously discussed, there are several experimental reasons why values may differ across these experiments [12]. Another recent paper tested RNA stacking [19], and we also see qualitative agreement between polarity pairs with the exception of G|A vs. A|G. Interestingly, we recently tested the effect of base stacking in stability of DNA tetrahedra [10] and we also found qualitative agreement in that study except with G|A and A|G. Due to the difference in experimental methods, it is difficult to say if this is a discrepancy or just reflective of the variables associated to each study. In both of these studies, stability was assessed by thermal melting rather than dissociation at constant temperature as in this study.

Overall, this data complements our previous work, completing quantification of the 16 possible base stacking energies in DNA. Differences in polarity turn out to be nontrivial, and our MD simulations give some insight into the origin of these differences. These types of stacking interactions have been shown experimentally to be important in assembly and stability of DNA structures [10,28], using fluorescent base analogs [29], and controlling enzymatic reactions [30], for example. It is our hope that the broader research community can use this work to inform design of sequences in these diverse applications.

