## Supplemental Information for "Investigating polarity effects in DNA base stacking"

**Contents**

**Table S1:** List of oligonucleotides

**Table S2:** Oligonucleotide combinations for each single-molecule construct

**Table S3**: Construct combination to form tethers with preferred base-stacking combination

**Figure S1**: Decay plot and single-exponential fitting of A|G, C|A, and C|G combinations of base-stacks at 15 pN

**Figure S2**: Decay plot and single-exponential fitting of T|A, T|G combinations and control construct at 15 pN

**Figure S3:** Decay plot and single-exponential fitting of T|C base-stack and corresponding control constructs at 15 pN

**Figure S4:** Polar plots for stacked bases in all MD simulations.

**Table S1.** List of oligonucleotide sequences.

| **Name** | **Sequence** | **Length** |
| --- | --- | --- |
| **Backbone sequences (5′-3′)**  **(Common set of oligos for all single-molecule constructs)** | | |
| 1. 5’Biotin | (5’ 2x bio) AACATCCAATAAATCATACAGGCAAGGCAAAGAATTAGCA | 40 |
| 2 | AAATTAAGCAATAAAGCCTC | 20 |
| 3 | AGAGCATAAAGCTAAATCGGTTGTACCAAAAACATTATGACCCTGTAATACTTTTGCGGG | 60 |
| 4 | AGAAGCCTTTATTTCAACGCAAGGATAAAAATTTTTAGAACCCTCATATATTTTAAATGC | 60 |
| 5 | AATGCCTGAGTAATGTGTAGGTAAAGATTCAAAAGGGTGAGAAAGGCCGGAGACAGTCAA | 60 |
| 6 | ATCACCATCAATATGATATTCAACCGTTCTAGCTGATAAATTAATGCCGGAGAGGGTAGC | 60 |
| 7 | TATTTTTGAGAGATCTACAAAGGCTATCAGGTCATTGCCTGAGAGTCTGGAGCAAACAAG | 60 |
| 8 | AGAATCGATGAACGGTAATCGTAAAACTAGCATGTCAATCATATGTACCCCGGTTGATAA | 60 |
| 9 | TCAGAAAAGCCCCAAAAACAGGAAGATTGTATAAGCAAATATTTAAATTGTAAACGTTAA | 60 |
| 10 | TATTTTGTTAAAATTCGCATTAAATTTTTGTTAAATCAGCTCATTTTTTAACCAATAGGA | 60 |
| 11 | ACGCCATCAAAAATAATTCGCGTCTGGCCTTCCTGTAGCCAGCTTTCATCAACATTAAAT | 60 |
| 12 | GTGAGCGAGTAACAACCCGTCGGATTCTCCGTGGGAACAAACGGCGGATTGACCGTAATG | 60 |
| 13 | GGATAGGTCACGTTGGTGTAGATGGGCGCATCGTAACCGTGCATCTGCCAGTTTGAGGGG | 60 |
| 14 | ACGACGACAGTATCGGCCTCAGGAAGATCGCACTCCAGCCAGCTTTCCGGCACCGCTTCT | 60 |
| 15 | GGTGCCGGAAACCAGGCAAAGCGCCATTCGCCATTCAGGCTGCGCAACTGTTGGGAAGGG | 60 |
| 16 | CGATCGGTGCGGGCCTCTTCGCTATTACGCCAGCTGGCGAAAGGGGGATGTGCTGCAAGG | 60 |
| 17 | CGATTAAGTTGGGTAACGCCAGGGTTTTCCCAGTCACGACGTTGTAAAACGACGGCCAGT | 60 |
| 18 | GCCAAGCTTGCATGCCTGCAGGTCGACTCTAGAGGATCCCCGGGTACCGAGCTCGAATTC | 60 |
| 19 | GTAATCATGGTCATAGCTGTTTCCTGTGTGAAATTGTTATCCGCTCACAATTCCACACAA | 60 |
| 20 | CATACGAGCCGGAAGCATAAAGTGTAAAGCCTGGGGTGCCTAATGAGTGAGCTAACTCAC | 60 |
| 21 | ATTAATTGCGTTGCGCTCACTGCCCGCTTTCCAGTCGGGAAACCTGTCGTGCCAGCTGCA | 60 |
| 22 | TTAATGAATCGGCCAACGCGCGGGGAGAGGCGGTTTGCGTATTGGGCGCCAGGGTGGTTT | 60 |
| 23 | TTCTTTTCACCAGTGAGACGGGCAACAGCTGATTGCCCTTCACCGCCTGGCCCTGAGAGA | 60 |
| 24 | GTTGCAGCAAGCGGTCCACGCTGGTTTGCCCCAGCAGGCGAAAATCCTGTTTGATGGTGG | 60 |
| 25 | TTCCGAAATCGGCAAAATCCCTTATAAATCAAAAGAATAGCCCGAGATAGGGTTGAGTGT | 60 |
| 26 | TGTTCCAGTTTGGAACAAGAGTCCACTATTAAAGAACGTGGACTCCAACGTCAAAGGGCG | 60 |
| 27 | AAAAACCGTCTATCAGGGCGATGGCCCACTACGTGAACCATCACCCAAATCAAGTTTTTT | 60 |
| 28 | GGGGTCGAGGTGCCGTAAAGCACTAAATCGGAACCCTAAAGGGAGCCCCCGATTTAGAGC | 60 |
| 29 | TTGACGGGGAAAGCCGGCGAACGTGGCGAGAAAGGAAGGGAAGAAAGCGAAAGGAGCGGG | 60 |
| 30 | CGCTAGGGCGCTGGCAAGTGTAGCGGTCACGCTGCGCGTAACCACCACACCCGCCGCGCT | 60 |
| 31 | TAATGCGCCGCTACAGGGCGCGTACTATGGTTGCTTTGACGAGCACGTATAACGTGCTTT | 60 |
| 32 | CCTCGTTAGAATCAGAGCGGGAGCTAAACAGGAGGCCGATTAAAGGGATTTTAGACAGGA | 60 |
| 33 | ACGGTACGCCAGAATCCTGAGAAGTGTTTTTATAATCAGTGAGGCCACCGAGTAAAAGAG | 60 |
| 34 | TCTGTCCATCACGCAAATTAACCGTTGTAGCAATACTTCTTTGATTAGTAATAACATCAC | 60 |
| 35 | TTGCCTGAGTAGAAGAACTCAAACTATCGGCCTTGCTGGTAATATCCAGAACAATATTAC | 60 |
| 36 | CGCCAGCCATTGCAACAGGAAAAACGCTCATGGAAATACCTACATTTTGACGCTCAATCG | 60 |
| 37 | TCTGAAATGGATTATTTACATTGGCAGATTCACCAGTCACACGACCAGTAATAAAAGGGA | 60 |
| 38 | CATTCTGGCCAACAGAGATAGAACCCTTCTGACCTGAAAGCGTAAGAATACGTGGCACAG | 60 |
| 39 | ACAATATTTTTGAATGGCTATTAGTCTTTAATGCGCGAACTGATAGCCCTAAAACATCGC | 60 |
| 40 | CATTAAAAATACCGAACGAACCACCAGCAGAAGATAAAACAGAGGTGAGGCGGTCAGTAT | 60 |
| 41 | TAACACCGCCTGCAACAGTGCCACGCTGAGAGCCAGCAGCAAATGAAAAATCTAAAGCAT | 60 |
| 42 | CACCTTGCTGAACCTCAAATATCAAACCCTCAATCAATATCTGGTCAGTTGGCAAATCAA | 60 |
| 43 | CAGTTGAAAGGAATTGAGGAAGGTTATCTAAAATATCTTTAGGAGCACTAACAACTAATA | 60 |
| 44 | GATTAGAGCCGTCAATAGATAATACATTTGAGGATTTAGAAGTATTAGACTTTACAAACA | 60 |
| 45 | ATTCGACAACTCGTATTAAATCCTTTGCCCGAACGTTATTAATTTTAAAAGTTTGAGTAA | 60 |
| 46 | CATTATCATTTTGCGGAACAAAGAAACCACCAGAAGGAGCGGAATTATCATCATATTCCT | 60 |
| 47 | GATTATCAGATGATGGCAATTCATCAATATAATCCTGATTGTTTGGATTATACTTCTGAA | 60 |
| 48 | TAATGGAAGGGTTAGAACCTACCATATCAAAATTATTTGCACGTAAAACAGAAATAAAGA | 60 |
| 49 | AATTGCGTAGATTTTCAGGTTTAACGTCAGATGAATATACAGTAACAGTACCTTTTACAT | 60 |
| 50 | CGGGAGAAACAATAACGGATTCGCCTGATTGCTTTGAATACCAAGTTACAAAATCGCGCA | 60 |
| 51 | GAGGCGAATTATTCATTTCAATTACCTGAGCAAAAGAAGATGATGAAACAAACATCAAGA | 60 |
| 52 | AAACAAAATTAATTACATTTAACAATTTCATTTGAATTACCTTTTTTAATGGAAACAGTA | 60 |
| 53 | CATAAATCAATATATGTGAGTGAATAACCTTGCTTCTGTAAATCGTCGCTATTAATTAAT | 60 |
| 54 | TTTCCCTTAGAATCCTTGAAAACATAGCGATAGCTTAGATTAAGACGCTGAGAAGAGTCA | 60 |
| 55 | ATAGTGAATTTATCAAAATCATAGGTCTGAGAGACTACCTTTTTAACCTCCGGCTTAGGT | 60 |
| 56 | TGGGTTATATAACTATATGTAAATGCTGATGCAAATCCAATCGCAAGACAAAGAACGCGA | 60 |
| 57 | GAAAACTTTTTCAAATATATTTTAGTTAATTTCATCTTCTGACCTAAATTTAATGGTTTG | 60 |
| 58 | AAATACCGACCGTGTGATAAATAAGGCGTTAAATAAGAATAAACACCGGAATCATAATTA | 60 |
| 59 | CTAGAAAAAGCCTGTTTAGTATCATATGCGTTATACAAATTCTTACCAGTATAAAGCCAA | 60 |
| 60 | CGCTCAACAGTAGGGCTTAATTGAGAATCGCCATATTTAACAACGCCAACATGTAATTTA | 60 |
| 61 | GGCAGAGGCATTTTCGAGCCAGTAATAAGAGAATATAAAGTACCGACAAAAGGTAAAGTA | 60 |
| 62 | ATTCTGTCCAGACGACGACAATAAACAACATGTTCAGCTAATGCAGAACGCGCCTGTTTA | 60 |
| 63 | TCAACAATAGATAAGTCCTGAACAAGAAAAATAATATCCCATCCTAATTTACGAGCATGT | 60 |
| 64 | AGAAACCAATCAATAATCGGCTGTCTTTCCTTATCATTCCAAGAACGGGTATTAAACCAA | 60 |
| 65 | GTACCGCACTCATCGAGAACAAGCAAGCCGTTTTTATTTTCATCGTAGGAATCATTACCG | 60 |
| 66 | CGCCCAATAGCAAGCAAATCAGATATAGAAGGCTTATCCGGTATTCTAAGAACGCGAGGC | 60 |
| 67 | GTTTTAGCGAACCTCCCGACTTGCGGGAGGTTTTGAAGCCTTAAATCAAGATTAGTTGCT | 60 |
| 68 | ATTTTGCACCCAGCTACAATTTTATCCTGAATCTTACCAACGCTAACGAGCGTCTTTCCA | 60 |
| 69 | GAGCCTAATTTGCCAGTTACAAAATAAACAGCCATATTATTTATCCCAATCCAAATAAGA | 60 |
| 70 | AACGATTTTTTGTTTAACGTCAAAAATGAAAATAGCAGCCTTTACAGAGAGAATAACATA | 60 |
| 71 | AAAACAGGGAAGCGCATTAGACGGGAGAATTAACTGAACACCCTGAACAAAGTCAGAGGG | 60 |
| 72 | TAATTGAGCGCTAATATCAGAGAGATAACCCACAAGAATTGAGTTAAGCCCAATAATAAG | 60 |
| 73 | AGCAAGAAACAATGAAATAGCAATAGCTATCTTACCGAAGCCCTTTTTAAGAAAAGTAAG | 60 |
| 74 | CAGATAGCCGAACAAAGTTACCAGAAGGAAACCGAGGAAACGCAATAATAACGGAATACC | 60 |
| 75 | CAAAAGAACTGGCATGATTAAGACTCCTTATTACGCAGTATGTTAGCAAACGTAGAAAAT | 60 |
| 76 | ACATACATAAAGGTGGCAACATATAAAAGAAACGCAAAGACACCACGGAATAAGTTTATT | 60 |
| 77 | TTGTCACAATCAATAGAAAATTCATATGGTTTACCAGCGCCAAAGACAAAAGGGCGACAT | 60 |
| 78 | TCAACCGATTGAGGGAGGGAAGGTAAATATTGACGGAAATTATTCATTAAAGGTGAATTA | 60 |
| 79 | TCACCGTCACCGACTTGAGCCATTTGGGAATTAGAGCCAGCAAAATCACCAGTAGCACCA | 60 |
| 80 | TTACCATTAGCAAGGCCGGAAACGTCACCAATGAAACCATCGATAGCAGCACCGTAATCA | 60 |
| 81 | GTAGCGACAGAATCAAGTTTGCCTTTAGCGTCAGACTGTAGCGCGTTTTCATCGGCATTT | 60 |
| 82 | TCGGTCATAGCCCCCTTATTAGCGTTTGCCATCTTTTCATAATCAAAATCACCGGAACCA | 60 |
| 83 | GAGCCACCACCGGAACCGCCTCCCTCAGAGCCGCCACCCTCAGAACCGCCACCCTCAGAG | 60 |
| 84 | CCACCACCCTCAGAGCCGCCACCAGAACCACCACCAGAGCCGCCGCCAGCATTGACAGGA | 60 |
| 85 | GGTTGAGGCAGGTCAGACGATTGGCCTTGATATTCACAAACAAATAAATCCTCATTAAAG | 60 |
| 86 | CCAGAATGGAAAGCGCAGTCTCTGAATTTACCGTTCCAGTAAGCGTCATACATGGCTTTT | 60 |
| 87 | GATGATACAGGAGTGTACTGGTAATAAGTTTTAACGGGGTCAGTGCCTTGAGTAACAGTG | 60 |
| 88 | CCCGTATAAACAGTTAATGCCCCCTGCCTATTTCGGAACCTATTATTCTGAAACATGAAA | 60 |
| 89 | GTATTAAGAGGCTGAGACTCCTCAAGAGAAGGATTAGGATTAGCGGGGTTTTGCTCAGTA | 60 |
| 90 | CCAGGCGGATAAGTGCCGTCGAGAGGGTTGATATAAGTATAGCCCGGAATAGGTGTATCA | 60 |
| 91 | CCGTACTCAGGAGGTTTAGTACCGCCACCCTCAGAACCGCCACCCTCAGAACCGCCACCC | 60 |
| 92 | TCAGAGCCACCACCCTCATTTTCAGGGATAGCAAGCCCAATAGGAACCCATGTACCGTAA | 60 |
| 93 | CACTGAGTTTCGTCACCAGTACAAACTACAACGCCTGTAGCATTCCACAGACAGCCCTCA | 60 |
| 94 | TAGTTAGCGTAACGATCTAAAGTTTTGTCGTCTTTCCAGACGTTAGTAAATGAATTTTCT | 60 |
| 95 | GTATGGGATTTTGCTAAACAACTTTCAACAGTTTCAGCGGAGTGAGAATAGAAAGGAACA | 60 |
| 96 | ACTAAAGGAATTGCGAATAATAATTTTTTCACGTTGAAAATCTCCAAAAAAAAGGCTCCA | 60 |
| 97 | AAAGGAGCCTTTAATTGTATCGGTTTATCAGCTTGCTTTCGAGGTGAATTTCTTAAACAG | 60 |
| 98 | CTTGATACCGATAGTTGCGCCGACAATGACAACAACCATCGCCCACGCATAACCGATATA | 60 |
| 99 | TTCGGTCGCTGAGGCTTGCAGGGAGTTAAAGGCCGCTTTTGCGGGATCGTCACCCTCAGC | 60 |
| 100 | AGCGAAAGACAGCATCGGAACGAGGGTAGCAACGGCTACAGAGGCTTTGAGGACTAAAGA | 60 |
| 101 | CTTTTTCATGAGGAAGTTTCCATTAAACGGGTAAAATACGTAATGCCACTACGAAGGCAC | 60 |
| 102 | CAACCTAAAACGAAAGAGGCAAAAGAATACACTAAAACACTCATCTTTGACCCCCAGCGA | 60 |
| 103 | TTATACCAAGCGCGAAACAAAGTACAACGGAGATTTGTATCATCGCCTGATAAATTGTGT | 60 |
| 104 | CGAAATCCGCGACCTGCTCCATGTTACTTAGCCGGAACGAGGCGCAGACGGTCAATCATA | 60 |
| 105 | AGGGAACCGAACTGACCAACTTTGAAAGAGGACAGATGAACGGTGTACAGACCAGGCGCA | 60 |
| 106 | TAGGCTGGCTGACCTTCATCAAGAGTAATCTTGACAAGAACCGGATATTCATTACCCAAA | 60 |
| 107 | TCAACGTAACAAAGCTGCTCATTCAGTGAATAAGGCTTGCCCTGACGAGAAACACCAGAA | 60 |
| 108 | CGAGTAGTAAATTGGGCTTGAGATGGTTTAATTTCAACTTTAATCATTGTGAATTACCTT | 60 |
| 109 | ATGCGATTTTAAGAACTGGCTCATTATACCAGTCAGGACGTTGGGAAGAAAAATCTACGT | 60 |
| 110 | TAATAAAACGAACTAACGGAACAACATTATTACAGGTAGAAAGATTCATCAGTTGAGATT | 60 |
| 111 | TAGGAATACCACATTCAACTAATGCAGATACATAACGCCAAAAGGAATTACGAGGCATAG | 60 |
| 112 | TAAGAGCAACACTATCATAACCCTCGTTTACCAGACGACGATAAAAACCAAAATAGCGAG | 60 |
| 113 | AGGCTTTTGCAAAAGAAGTTTTGCCAGAGGGGGTAATAGTAAAATGTTTAGACTGGATAG | 60 |
| 114 | CGTCCAATACTGCGGAATCGTCATAAATATTCATTGAATCCCCCTCAAATGCTTTAAACA | 60 |
| 115 | GTTCAGAAAACGAGAATGACCATAAATCAAAAATCAGGTCTTTACCCTGACTATTATAGT | 60 |
| 116 | CAGAAGCAAAGCGGATTGCATCAAAAAGATTAAGAGGAAGCCCGAAAGACTTCAAATATC | 60 |
| 117 | GCGTTTTAATTCGAGCTTCAAAGCGAACCAGACCGGAAGCAAACTCCAACAGGTCAGGAT | 60 |
| 118 | TAGAGAGTACCTTTAATTGCTCCTTTTGATAAGAGGTCATTTTTGCGGATGGCTTAGAGC | 60 |
| 119 | TTAATTGCTGAATATAATGCTGTAGCTCAACATGTTTTAAATATGCAACTAAAGTACGGT | 60 |
| 120 | GTCTGGAAGTTTCATTCCATATAACAGTTGATTCCCAATTCTGCGAACGAGTAGATTTAG | 60 |
| 121 | TTTGACCATTAGATACATTTCGCAAATGGTCAATAACCTGTTTAGCTAT | 49 |
| 122 | ATTTTCATTTGGGGCGCGAGCTGAAAAGGT | 30 |
| CutOligo | CTACTAATAGTAGTAGCATTAACATCCAATAAATCATACA | 40 |
| **Sequence used in specific combination for each construct (5′-3′)**  **Overhanging regions underlined, spacer T are marked in blue, and stacking bases in red** | | |
| OH-A | GGCATCAATTCTACTAATAGTAGTAGCATTCCGTGCCTGTGAACGAGCTGCCCCATGGCA | 60 |
| OH-G | GGCATCAATTCTACTAATAGTAGTAGCATTCCGTGCCTGTGAACGAGCTGCCCCATGGCG | 60 |
| OH-C | GGCATCAATTCTACTAATAGTAGTAGCATTCCGTGCCTGTGAACGAGCTGCCCCATGGCC | 60 |
| OH-T | GGCATCAATTCTACTAATAGTAGTAGCATTCCGTGCCTGTGAACGAGCTGCCCCATGGCT | 60 |
| A:C-A | ACGTCGCCTGCCATGGGGCAGCTCGTTCACAGGCACGG | 38 |
| T:T-G | GGCGACGTAGCCATGGGGCAGCTCGTTCACAGGCACGG | 38 |
| T:C-A | ACGTCGCCAGCCATGGGGCAGCTCGTTCACAGGCACGG | 38 |
| C:T-G | GGCGACGTGGCCATGGGGCAGCTCGTTCACAGGCACGG | 38 |
| C:C-A | ACGTCGCCGGCCATGGGGCAGCTCGTTCACAGGCACGG | 38 |
| T:G-T | TGCAGCGGAGCCATGGGGCAGCTCGTTCACAGGCACGG | 38 |
| T: Sp-T-G | GGCGACGTTTTAGCCATGGGGCAGCTCGTTCACAGGCACGG | 41 |
| T: Sp-C-A | ACGTCGCCTTTAGCCATGGGGCAGCTCGTTCACAGGCACGG | 41 |
| T: Sp-A-C | CCGCTGCATTTAGCCATGGGGCAGCTCGTTCACAGGCACGG | 41 |
| T: Sp-G-T | TGCAGCGGTTTAGCCATGGGGCAGCTCGTTCACAGGCACGG | 41 |

**Table S2:** Oligonucleotide combinations for each single-molecule construct.

| **Construct #** | **Oligo Mix (1:15:50 mole ratio)** |
| --- | --- |
| 1 | Oligos 1-122, OH-A, A:C-A |
| 2 | Oligos 1-122, OH-T, T:T-G |
| 3 | Oligos 1-122, OH-T, T:C-A |
| 4 | Oligos 1-122, OH-C, C:T-G |
| 5 | Oligos 1-122, OH-C, C:C-A |
| 6 | Oligos 1-122, OH-T, T:G-T |
| 7 | Oligos 1-122, OH-T, T:Sp-T-G |
| 8 | Oligos 1-122, OH-T, T:Sp-C-A |
| 9 | Oligos 1-122, OH-T, T:Sp-A-C |
| 10 | Oligos 1-122, OH-T, T:Sp-G-T |

**Table S3:** Construct combination to form tethers in single-molecule experiments**.**

| **Tether ID** | **Stacking Combination (5′\|3′)** | **Construct combinations to form tether** |
| --- | --- | --- |
| 1 | A\|G | 1& 7 |
| 2 | T\|A | 2 & 8 |
| 3 | T\|G | 3 & 7 |
| 4 | C\|A | 4 & 8 |
| 5 | C\|G | 5 & 7 |
| 8 | Control for (A\|G, T\|A, T\|G, C\|A, C\|G) | 7 & 8 |
| 9 | T\|C | 6 & 9 |
| 12 | Control for (T\|C) | 9 &10 |

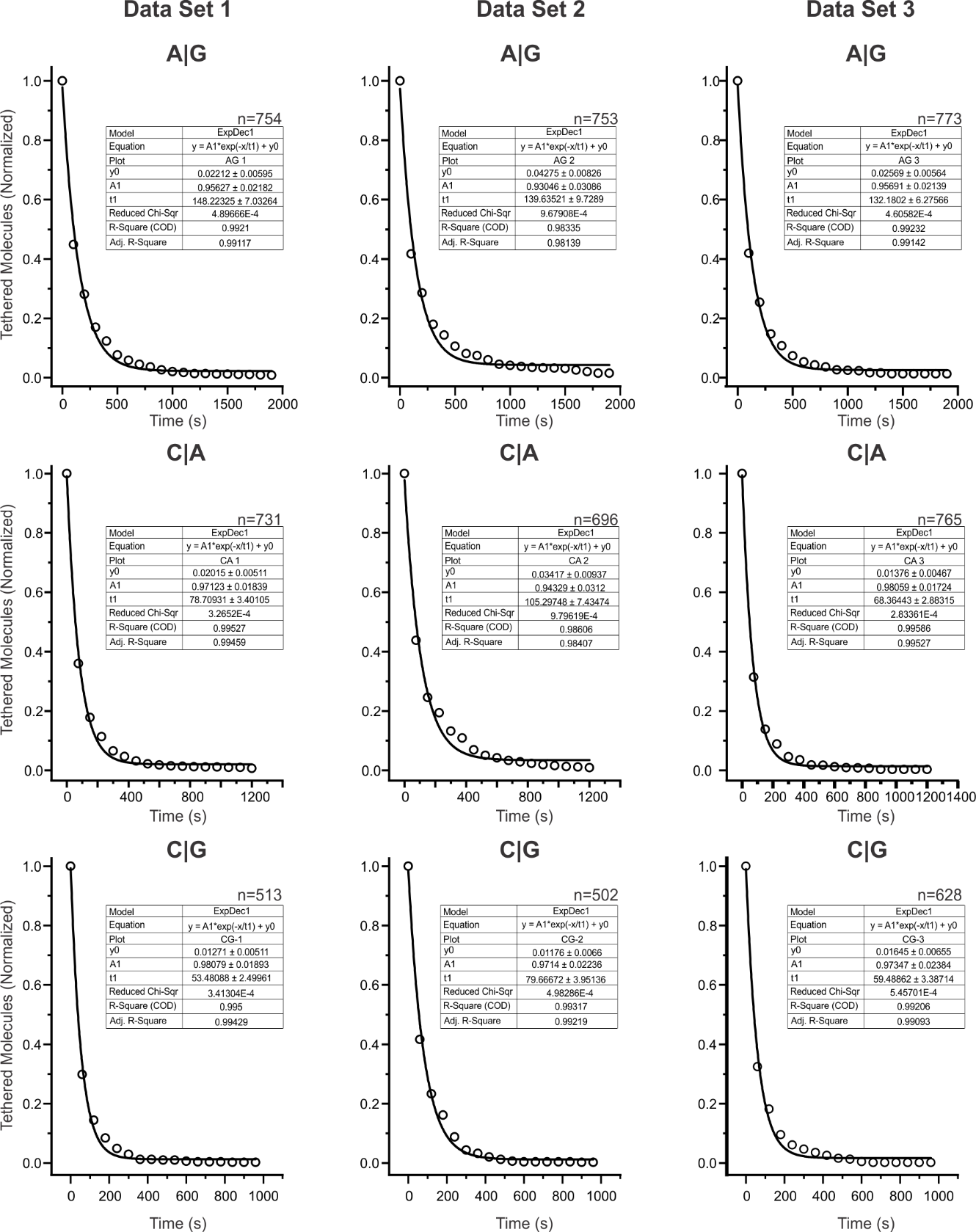

**Figure S1**: Decay plot and single-exponential fitting of A|G, C|A, and C|G combinations of base-stacks at 15 pN.

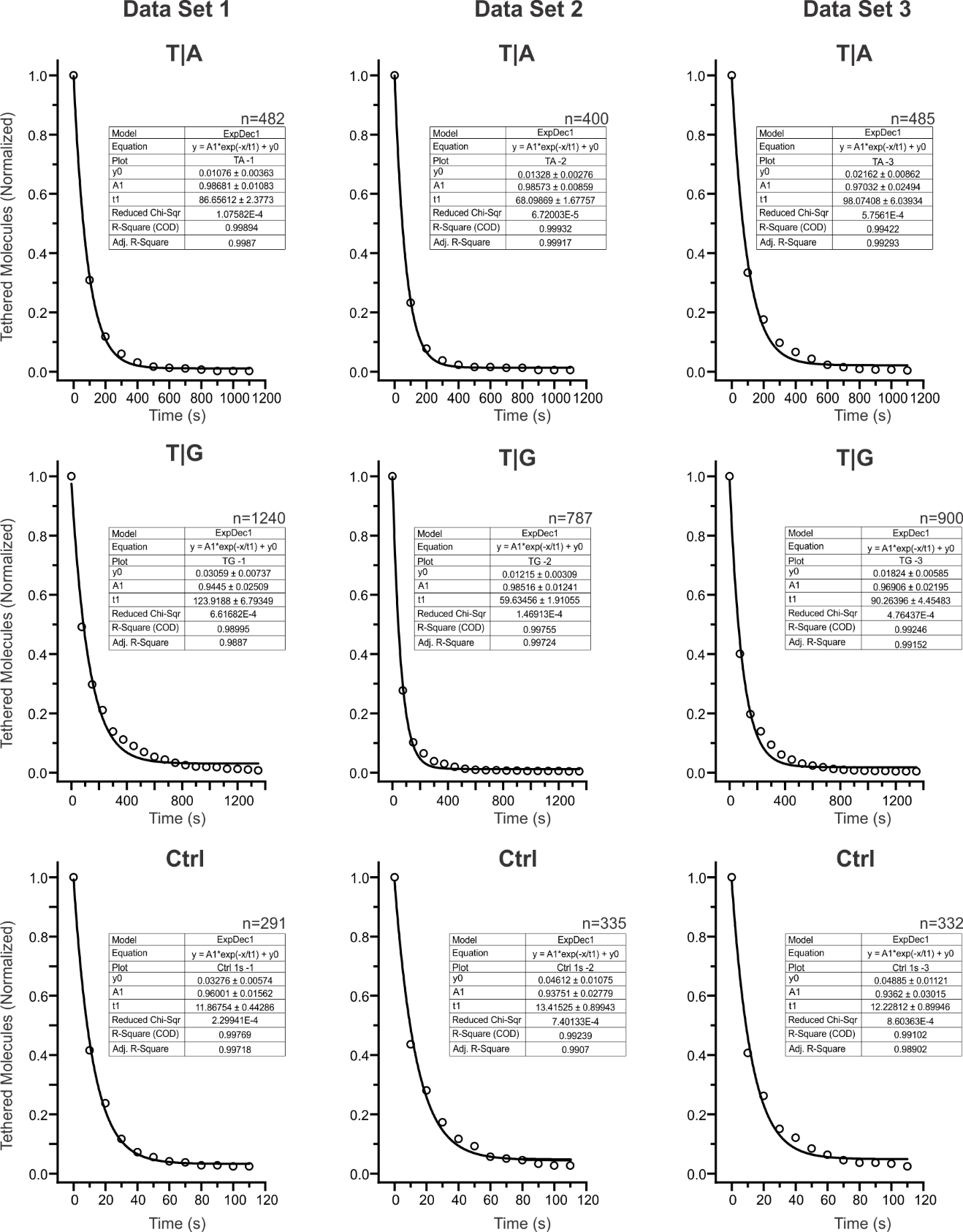

**Figure S2:** Decay plot and single-exponential fitting of T|A, T|G combinations and control construct at 15 pN.

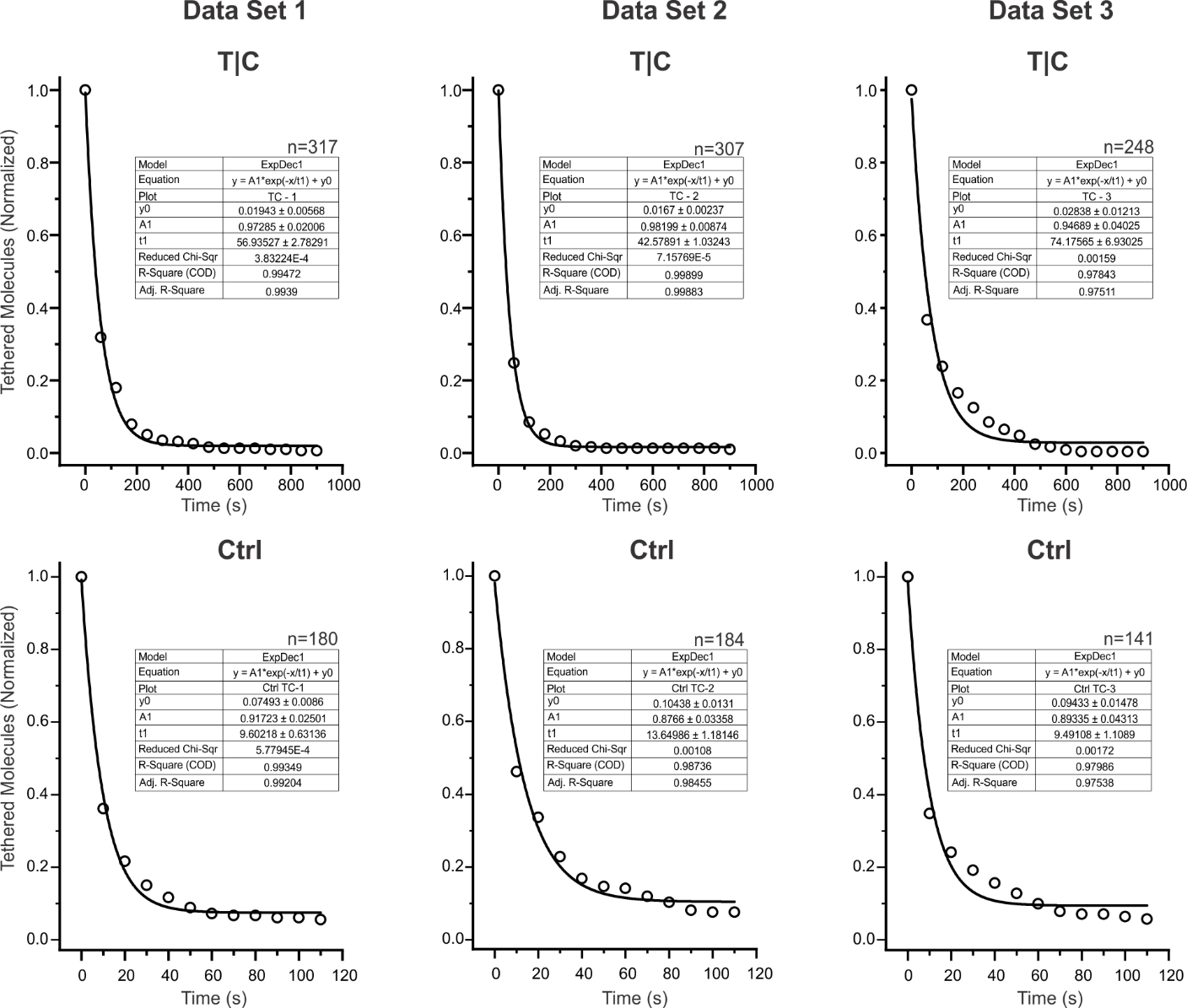

**Figure S3:** Decay plot and single-exponential fitting of T|C base-stack and corresponding control constructs at 15 pN.

|  | 300 K | 320 K | 340 K | 360 K | 380 K | 400 K |
| --- | --- | --- | --- | --- | --- | --- |
| A\|C | 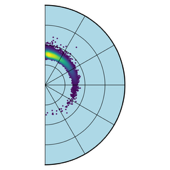 | 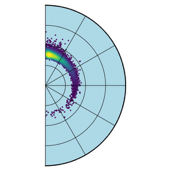 | 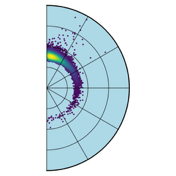 | 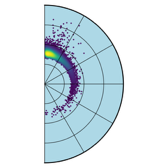 | 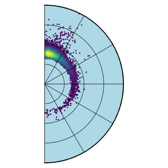 | 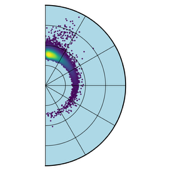 |
| C\|A | 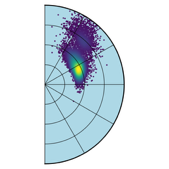 | 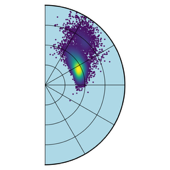 | 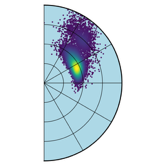 | 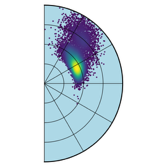 | 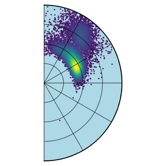 | 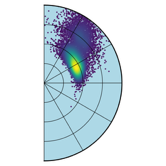 |
| A\|G | 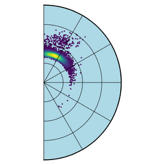 | 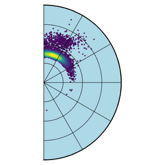 | 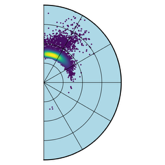 | 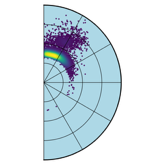 | 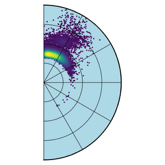 | 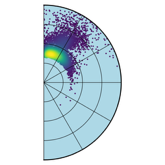 |
| G\|A | 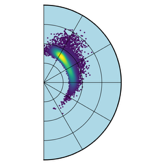 | 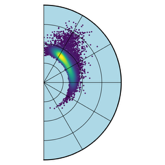 | 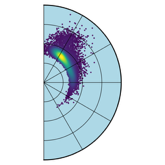 | 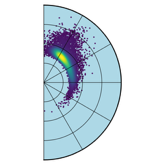 | 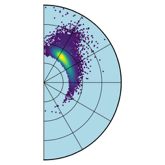 | 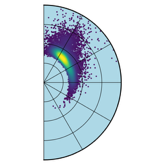 |
| A\|T | 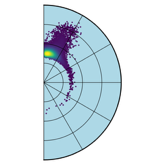 | 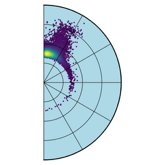 | 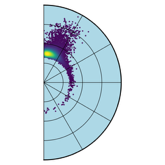 | 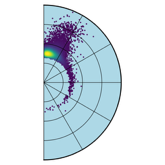 | 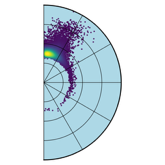 | 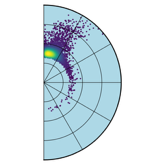 |
| T\|A | 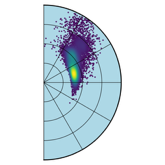 | 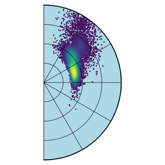 | 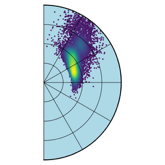 | 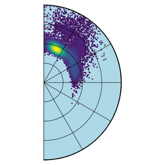 | 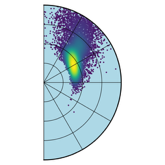 | 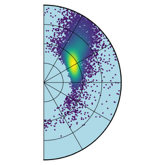 |
| C\|G | 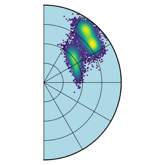 | 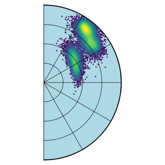 | 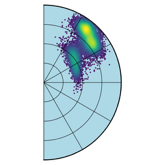 | 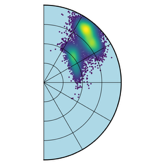 | 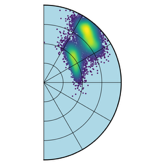 | 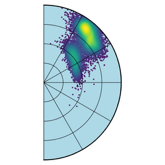 |
| G\|C | 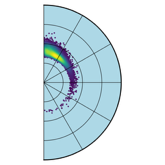 | 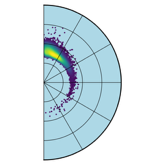 | 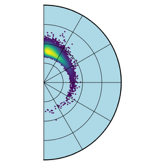 | 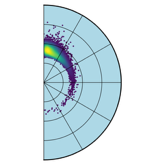 | 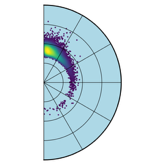 | 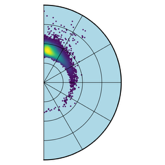 |
| C\|T | 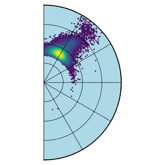 | 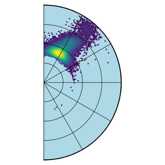 | 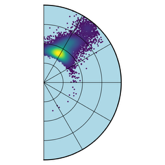 | 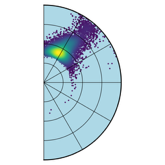 | 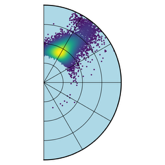 | 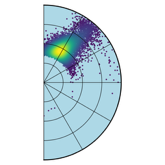 |
| T\|C | 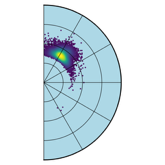 | 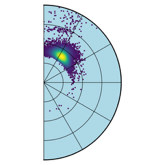 | 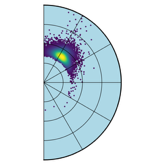 | 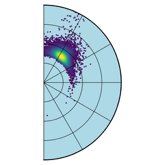 | 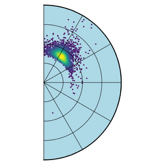 | 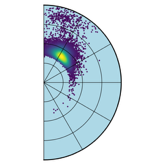 |
| G\|T | 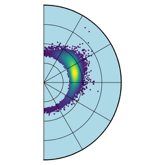 | 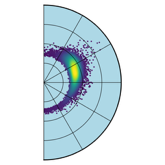 | 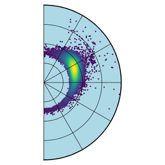 | 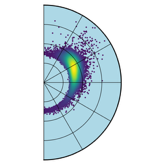 | 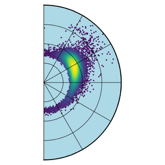 | 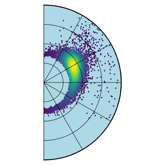 |
| T\|G | 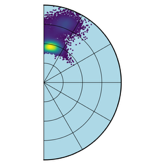 | 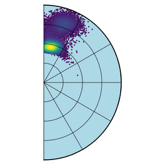 | 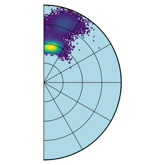 | 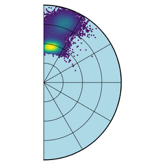 | 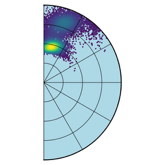 | 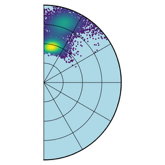 |

**Figure S4:** Polar plots for stacked bases in all MD simulations.
